# Killing of Gram-negative and Gram-positive bacteria by a bifunctional cell wall-targeting T6SS effector

**DOI:** 10.1101/2021.03.04.433973

**Authors:** Nguyen-Hung Le, Victor Pinedo, Juvenal Lopez, Felipe Cava, Mario Feldman

## Abstract

The type VI secretion system (T6SS) is a powerful tool deployed by Gram-negative bacteria to antagonize neighboring organisms. Here, we report that *Acinetobacter baumannii* ATCC 17978 (Ab17978) secretes D-lysine (D-Lys), increasing the extracellular pH and enhancing the peptidoglycanase activity of the T6SS effector Tse4. This synergistic effect of D-Lys on Tse4 activity enables Ab17978 to outcompete Gram-negative bacterial competitors, demonstrating that bacteria can modify their microenvironment to increase their fitness during bacterial warfare. Remarkably, this lethal combination also results in T6SS-mediated killing of Gram-positive bacteria. Further characterization revealed that Tse4 is a bifunctional enzyme consisting of both lytic transglycosylase and endopeptidase activities, thus representing a novel family of modularly organized T6SS PG degrading effectors with an unprecedented impact in antagonistic bacterial interactions.

**One sentence summary:** By modulating local environmental pH through D-Lys secretion, *Acinetobacter baumannii* enhances the activity of a bifunctional cell wall-targeting T6SS effector, increasing its killing activity against Gram-negative and -positive competitors.

## Main text

Bacteria live in dense communities and are often in constant competition with other bacterial species to secure nutrients and space. Bacterial warfare is mediated by the production of diffusible antimicrobial compounds and by sophisticated molecular weapons, such as the type VI secretion system (T6SS). The T6SS is a dynamic nanomachine that delivers toxic effector proteins from an attacking cell (predator) to nearby competitors (prey) in a contact-dependent manner. Although bacteria possess a diverse arsenal of toxic effector proteins to kill Gram-negative bacteria, current evidence suggests that Gram-positive bacteria are immune to T6SS attacks. Immunity to T6SS-dependent killing between Gram-negative kin cells is accomplished by the expression of immunity proteins, which specifically bind and inactivate their cognate effector. Broad-spectrum mechanisms of protection against non-kin T6SS attacks in Gram-negative bacteria have only recently been uncovered(*1*, *2*).

Due to its essentiality, the bacterial cell wall peptidoglycan (PG), also known as murein, is targeted by various T6SS effectors. PG is composed of glycan chains of alternating N-acetylglucosamine (GlcNAc) and N-acetylmuramic acid (MurNAc) that are crosslinked through MurNAc-attached peptides. PG-degrading (PGase) effectors induce bacterial cell lysis, which facilitates the complete clearance of bacterial competitors by preventing dead cells from shielding susceptible prey from T6SS attacks, a phenomenon known as the “corpse barrier” effect(*3*). The PGase effectors characterized to date contain one lytic enzymatic activity and can be categorized as muramidases, which cleave the β-(1,4)-glycosidic bond(*4*, *5*), N-acetylmuramyl-L-alanine amidases, which unlink the peptides chains from MurNAc (*6*) and LD-(*4*, *7*), or DD-endopeptidases(*7*), which cleave PG crosslinks.

T6SS-dependent bacterial warfare generates an arms race in which both predator and prey evolve novel tools to prevail. We recently showed that some *Acinetobacter baumannii* strains, such as Ab17978, use the periplasmic racemase RacK to produce the non-canonical D-amino acid (NCDAA) D-lysine (D-Lys). Incorporation of D-Lys into the PG of Ab17978 underlies a defensive strategy against the PGase activity of T6SS effectors from competing bacteria(*2*). Beyond being incorporated into the PG, most of the D-Lys that is produced is secreted into the extracellular milieu, where it accumulates to millimolar concentrations(*2*). It has been recently suggested that NCDAAs carry out diverse biological roles in bacterial ecosystems(*8*). Therefore, we decided to investigate the role of extracellular D-Lys in bacterial warfare.

Previous work has shown that T6SS-dependent toxicity is impacted by the extracellular environment(*9*). We hypothesized that D-Lys secretion could potentiate T6SS-dependent bacterial killing by Ab17978. To test this hypothesis, we compared the T6SS killing activity of wild-type (WT) Ab17978 with that of its *racK* deletion (*ΔracK*) derivative. We found that the T6SS of WT Ab17978 is ~100 fold more lethal than the *ΔracK* strain against two different prey, *Escherichia coli* MG1655 and *Acinetobacter nosocomialis* M2 with an inactive T6SS (M2*ΔtssB*) (Fig. 1A). As expected, Ab17978*ΔtssM*, which possesses an inactive T6SS, did not display bacterial killing (Fig. 1A). WT and *ΔracK* secreted equal amounts of the T6SS protein Hcp, indicating that deletion of *racK* does not affect T6SS dynamics (Fig. S1). We also confirmed that Ab17978ΔtssM secretes similar amounts of D-Lys compared with the WT strain, suggesting that T6SS functionality and D-Lys secretion are independent processes (Fig. S2). Remarkably, heterologous *racK* expression in the clinical isolate *A. baumannii* strain UPAB1, which does not encode *racK*, resulted in a 100-fold increase in T6SS lethality compared with WT (Fig. 1B). Together, these data demonstrate that D-Lys secretion enhances the T6SS-mediated killing of *A. baumannii*.

**Fig. 1:**
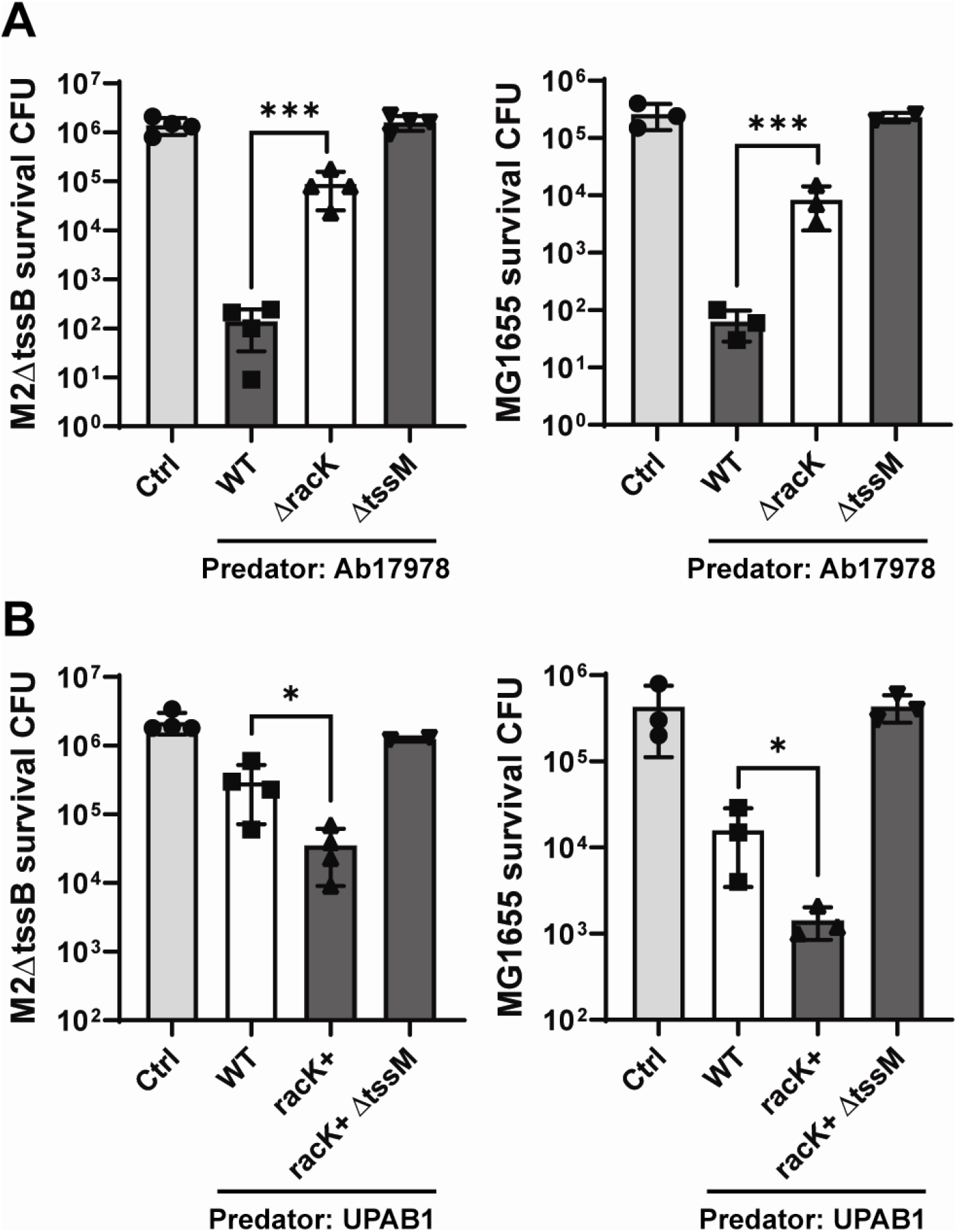
RacK enhances T6SS-dependent killing by *A. baumannii*. **A.** Competition assay using Ab17978 WT, *ΔracK*, or *ΔtssM* as predators and *A. nocosomialis* M2*ΔtssB* or *E. coli* MG1655 as prey. **B.** Competition assay using *A. baumannii* UPAB1 WT, UPAB1 heterologously expressing *racK* (racK+) or racK+*ΔtssM* as predators and *A. nocosomialis* M2*ΔtssB* or *E. coli* MG1655 as prey. Bar graphs represent means of prey survival CFU after 4 h of co-incubation ± SD of at least three biological replicates. Statistical analyses were performed using the unpaired Student’s *t* test, \**p* < 0.05 and \*\*\**p* < 0.001.

Ab17978 encodes four T6SS effectors: Tse1 is a predicted lipase; Tse2 is a predicted nuclease; Tse3 is an effector of unknown function; and Tse4 (ACX60_00605) is predicted to contain PGase activity (*10*). To gain insight into the effectors involved in D-Lys-mediated enhancement of T6SS lethality, we tested the killing efficiency of mutants unable to secrete one or more effectors. We found that a mutant strain unable to secrete effectors Tse1, Tse2 and Tse3 (*Δ123*) retained most of its killing ability against M2*ΔtssB*, indicating that Tse4 plays a major role in bacterial killing (Fig. 2). A *ΔracK* derivative of the *Δ123* strain (*Δ123ΔracK*), lost most of its killing capacity, indicating that RacK enhances Tse4 toxicity. Immunity proteins are generally encoded adjacently to their cognate effector. Indeed, overexpression of *tsi4* (ACX60_00610) in M2*ΔtssB* prey prevented killing by Ab17978*Δ123*, indicating that Tsi4 is the immunity protein of Tse4 (Fig. 2). Furthermore, the enhanced killing provided by RacK disappeared when M2*ΔtssB* expressed *tsi4* (Fig. 2). Together, our results strongly suggest that the synergistic effect of RacK on T6SS lethality is linked to the cell wall-targeting effector Tse4.

**Fig. 2:**
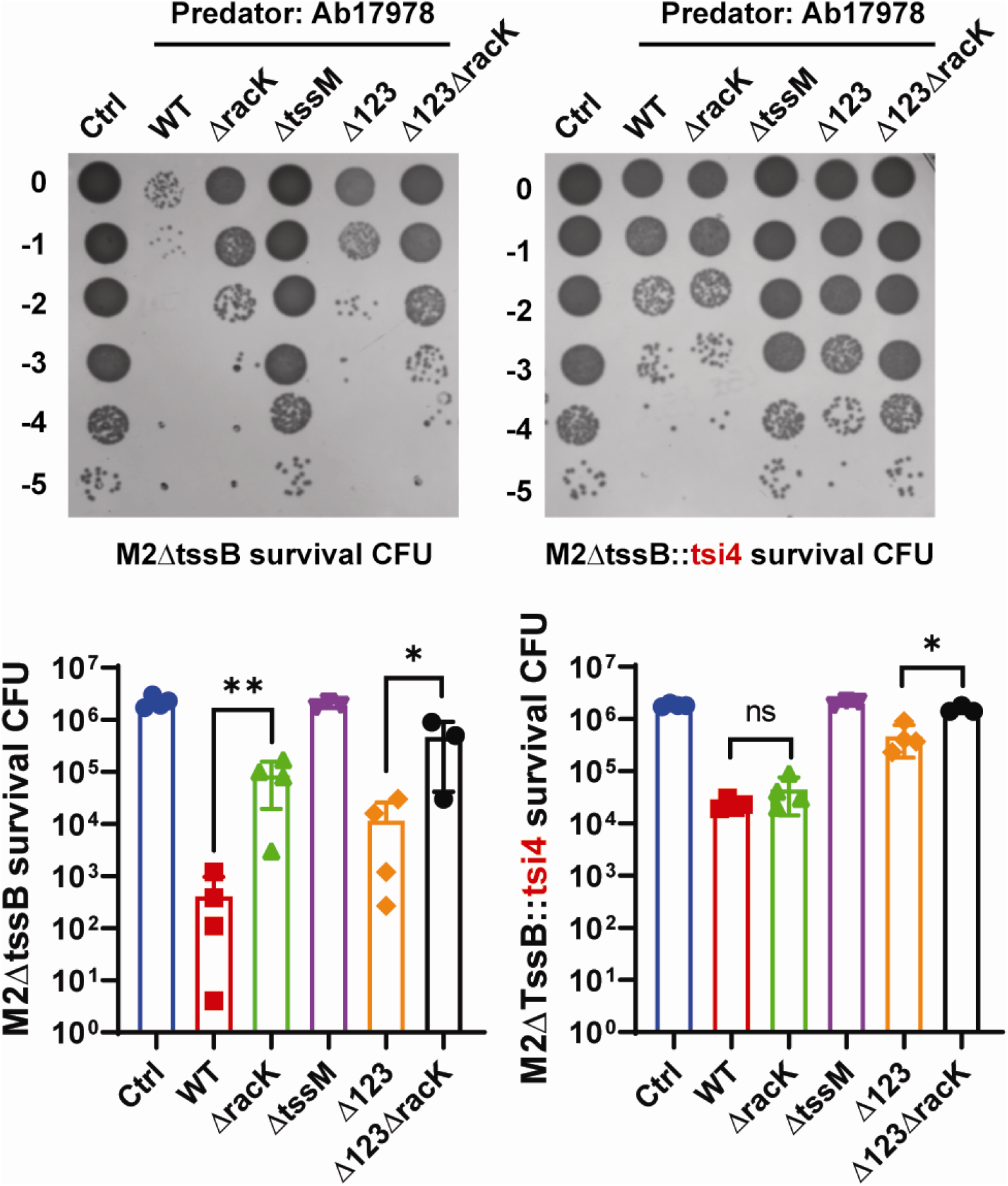
Synergistic effect of RacK on bacterial killing is linked to PG hydrolase effector Tse4. Competition assay using Ab17978 WT, *ΔracK*, *ΔtssM*, *Δ123*, *Δ123ΔracK* as predators and M2*ΔtssB* either harboring an empty vector (left panel) or overexpressing Tsi4, the immunity protein of Tse4 (right panel) as prey. Bar graphs represent means of prey survival CFU after 4 h of co-incubation ± SD of at least three biological replicates. Statistical analyses were performed using the unpaired Student’s *t* test, \**p* < 0.05, \*\**p* < 0.01 and *ns* not significant.

Next, we performed interbacterial competition assays between the Ab17978*ΔracK* predator and M2*ΔtssB* prey in media supplemented with increasing amounts of D-Lys. Consistent with our previous result, we found that T6SS-dependent prey killing was augmented with increasing concentrations of D-Lys. The effect was noticeable at 5 mM D-Lys and led to WT killing levels at 20 mM D-Lys (Fig. S3A). Importantly, growth of Ab17978 predator strains and M2 prey strains was not affected by the presence of 20 mM D-Lys in the media (Fig. S5), thus excluding the possible toxicity of D-Lys alone. Extracellular NCDAAs, such as D-Met, can be incorporated into the PG of non-producing species(*11*). To test whether the enhanced T6SS lethality observed in the presence of D-Lys can be emulated by other NCDAAs, we performed bacterial killing assays in the presence of D-Met, D-Ala, or D-Arg. While D-Met and D-Ala did not potentiate T6SS lethality (Fig. S3C, D), D-Arg mimicked the effect of D-Lys (Fig. S3B). Unexpectedly, we found that L-Lys and L-Arg also synergized with the T6SS (Fig. S3E). These results indicate that basic amino acids, in either their L- or D- form, enhance the lethality of the T6SS. Thus, we then hypothesized that the RacK-dependent enhancement of interbacterial killing was due to a D-Lys-mediated increase in the pH at the interface between WT Ab17978 and its prey. Supporting this concept, when killing assays were performed in a buffered pH of 6.8 (the pH of LB media) we found that the presence of RacK or 20mM D-Lys no longer increased the bactericidal effect of Ab17978 (Fig. 3A, Fig. S4). In contrast, bacterial killing by Ab17978 WT and *ΔracK* was increased in a buffered pH of 8.0 (Fig. 3A). We hypothesized that the increased T6SS-dependent killing at alkaline pH could be due to enhanced Tse4 activity in this condition. To this end, we tested the *in vitro* activity of purified Tse4 using a Remazol Brilliant Blue (RBB)-dye release assay and found that the enzymatic activity of Tse4 was optimal at pH 8 and above (Fig. 3B). Based on this data, we propose a model in which D-Lys secretion increases T6SS-mediated killing by creating an alkaline microenvironment in the predator-prey interface, resulting in maximal Tse4 activity.

**Fig. 3:**
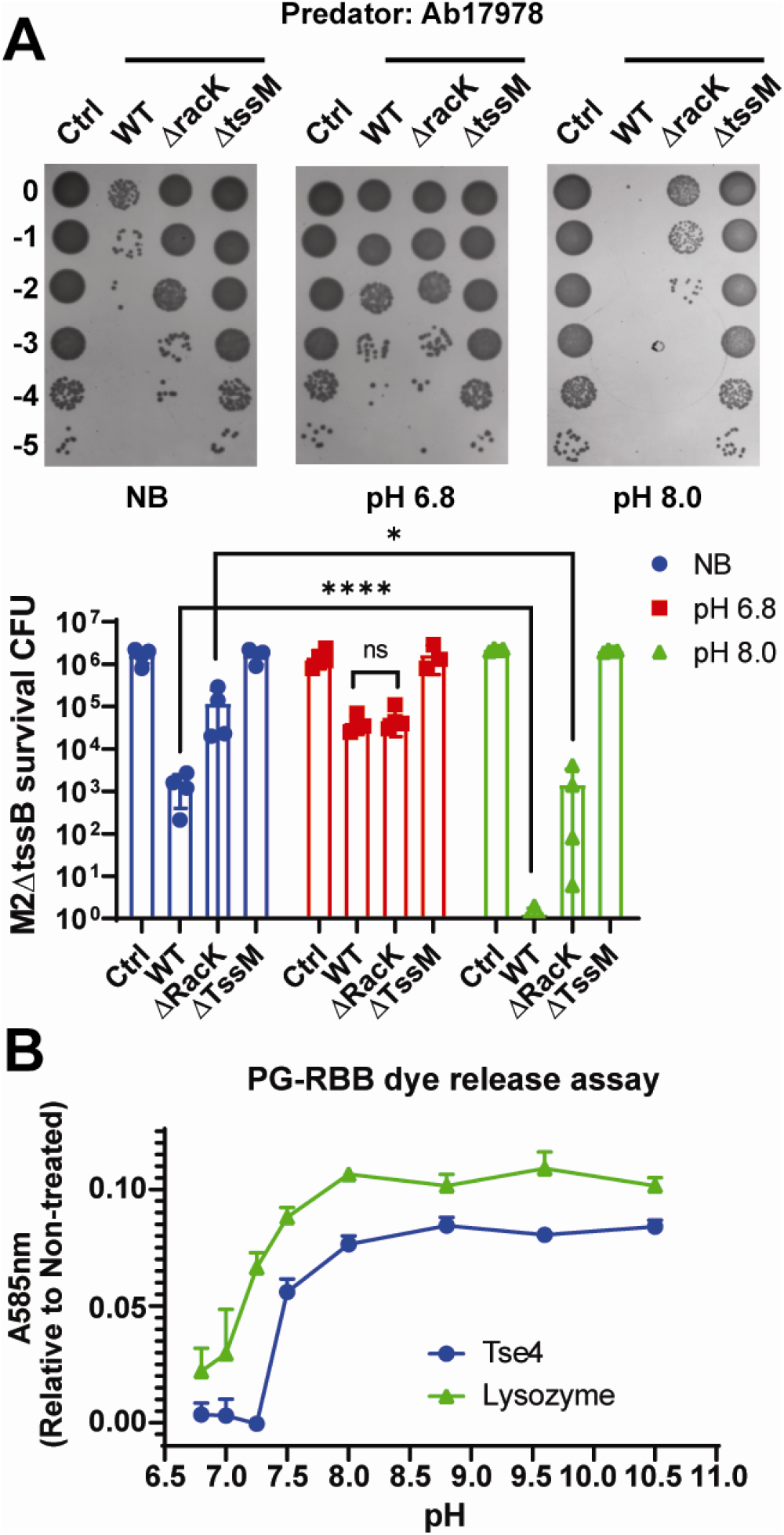
Alkaline environmental conditions enhance bacterial killing and Tse4 activity *in vitro*. **A.** Competition assay using Ab17978 WT, *ΔracK*, or *ΔtssM* as predators and M2*ΔtssB* as prey in non-buffered (NB) media or media buffered at pH 6.8 or 8.0. Bar graphs represent means of prey survival CFU after 4 h of co-incubation of four biological replicates. **B.** Remazol Brilliant Blue (RBB)-labeled sacculi were incubated with 7.5 μM lysozyme or purified Tse4. After ~16 h, the undigested PG was pelleted and released dye was quantified by measuring dye absorbance at 585 nm. Values represent mean of 3 independent enzymatic assays ± SD.

The dramatic RacK-induced killing efficiency of Ab17978 prompted us to test if this bacterium is able to kill Gram-positive bacteria. Remarkably, we found that Ab17978 can kill strains of *Bacillus subtilis*, *Listeria monocytogenes*, and methicillin-resistant *Staphylococcus aureus* (MRSA) in a T6SS- and RacK-dependent manner (Fig. 4A). These Gram-positive prey did not inhibit Ab17978 growth during co-incubation (Fig. S6). Notably, we found that the *Δ4* strain, which lacks *tse4*, did not kill any of these strains, while the *Δ123* strain exhibited increased killing in all cases. We hypothesize that the *Δ123* strain secretes increased amounts of Tse4 due to the lack of competition for the T6SS machinery, which is consistent with previous work in *Pseudomonas aeruginosa*(*12*). Importantly, addition of purified Tse4 to *B. subtilis* cultures did not induce cell lysis (Fig. 4B), and co-incubation of Ab17978 and *B. subtilis* in liquid media did not result in bacterial killing (Fig. S7), indicating that T6SS-dependent activity against Gram-positive bacteria is contact-dependent. Our results are the first demonstration of T6SS-dependent killing of Gram-positive bacteria.

**Fig. 4:**
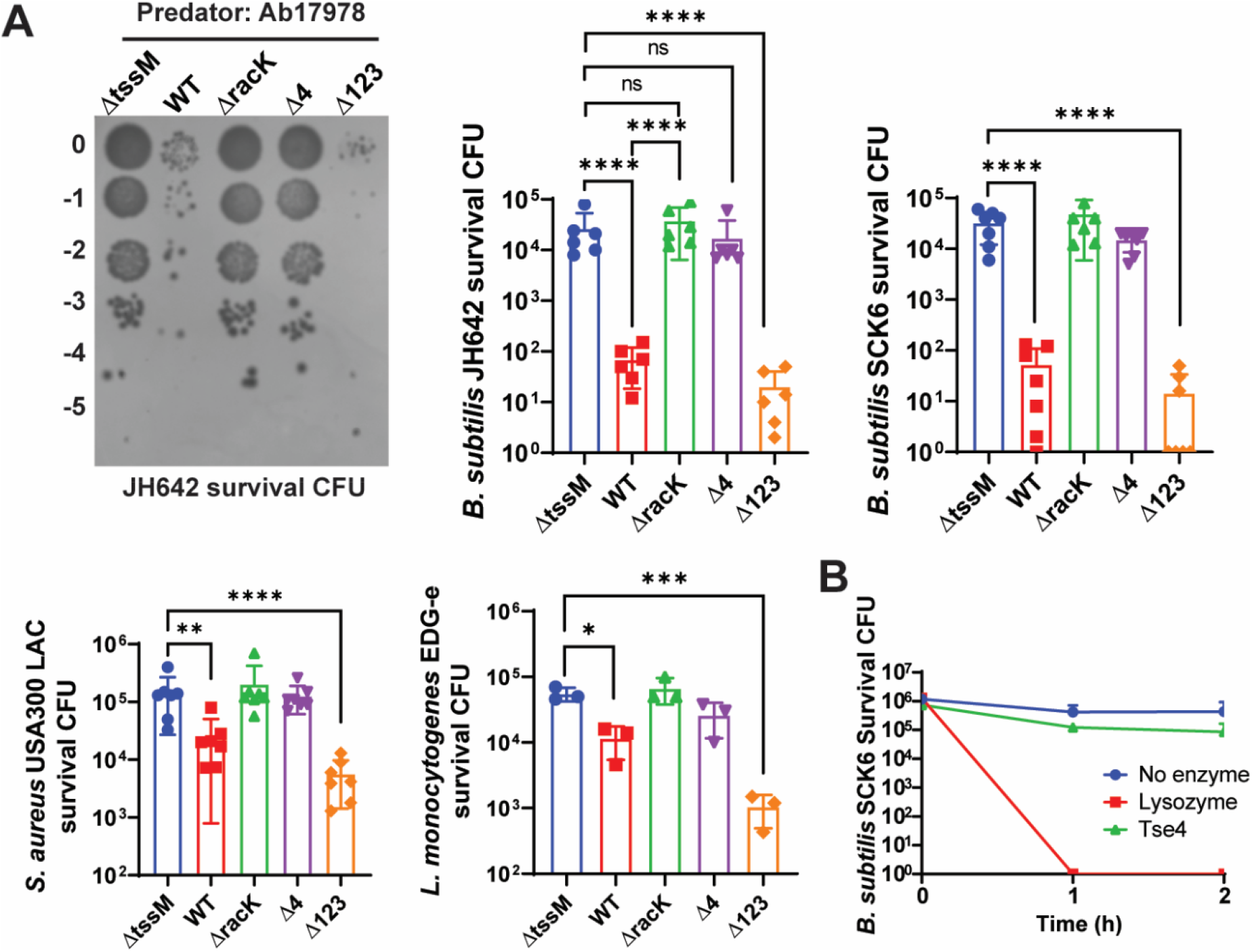
Ab17978 kills various Gram-positive bacteria in a Tse4- and contact-dependent manner. **A.** Competition assay using Ab17978 WT, *ΔtssM*, *ΔracK*, *Δ4* or *Δ123* as predators and *B. subtilis* JH642 or SCK6, *S. aureus* USA300 LAC, and *L. monocytogenes* EDG-e as prey. Bar graphs represent means of prey survival CFU after co-incubation ± SD of at least three biological replicates. Statistical analyses were performed using the unpaired Student’s *t* test, \**p* < 0.05, \*\**p* < 0.01, ***p < 0.001, ****p < 0.0001 and ns, not significant. **B.** Late exponential phase growing *B. subtilis* cells (OD600 ~ 1) were treated with either 7.5 μM of lysozyme or Tse4 for 2 h. Survival CFUs were enumerated at 0, 1 and 2 h of treatment. Values represent means ± SD of three biological replicates.

As for *E. coli,* PG of *B. subtilis* and *L. monocytogenes* use D-Ala-meso diaminopimelate (mDAP) crosslinks (Fig. 5A)(*13*). In contrast, *S. aureus* PG uses a penta-Gly bridge connecting the peptide stems (Fig. 5A)(*13*). Therefore, Tse4 has the remarkable ability to degrade different PG chemotypes. The Conserved Domain Architecture Retrieval Tool (CDART) algorithm predicted that Tse4 has three conserved domains: a LysM domain in the N-terminal region, followed by a lysozyme-like domain and a Zinc-binding peptidase domain. Consistently, the Phyre2 server identified two conserved catalytic domains in Tse4 (Fig. S8). Amino acid residues 242-447 modeled to a lysozyme-like domain structurally similar to lytic transglycosylase gp144 of bacteriophage phiKZ (PDB: 3BKH), and this domain would therefore be predicted to non-hydrolytically cleave the glycan chains between GlcNAc and MurNAc, thus producing anhydromuropeptides. Amino acid residues 634-798 modeled to two distinct structures. In the first, the peptidase domain was modeled to the DD-endopeptidase ShyA of *Vibrio cholerae* (PDB:6U2A), which cleaves D-Ala-mDAP bonds(*14*). In the second, the peptidase domain was modeled to the lysostaphin LytM of *S. aureus* (PDB: 2B44), an antibacterial enzyme that is capable of cleaving the penta-Gly bridges found in the PG crosslinks of staphylococci(*15*). Thus, this analysis suggests that Tse4 is likely a bifunctional (lytic transglycosylase and endopeptidase) effector capable of targeting the cell wall of various bacteria.

**Fig. 5:**
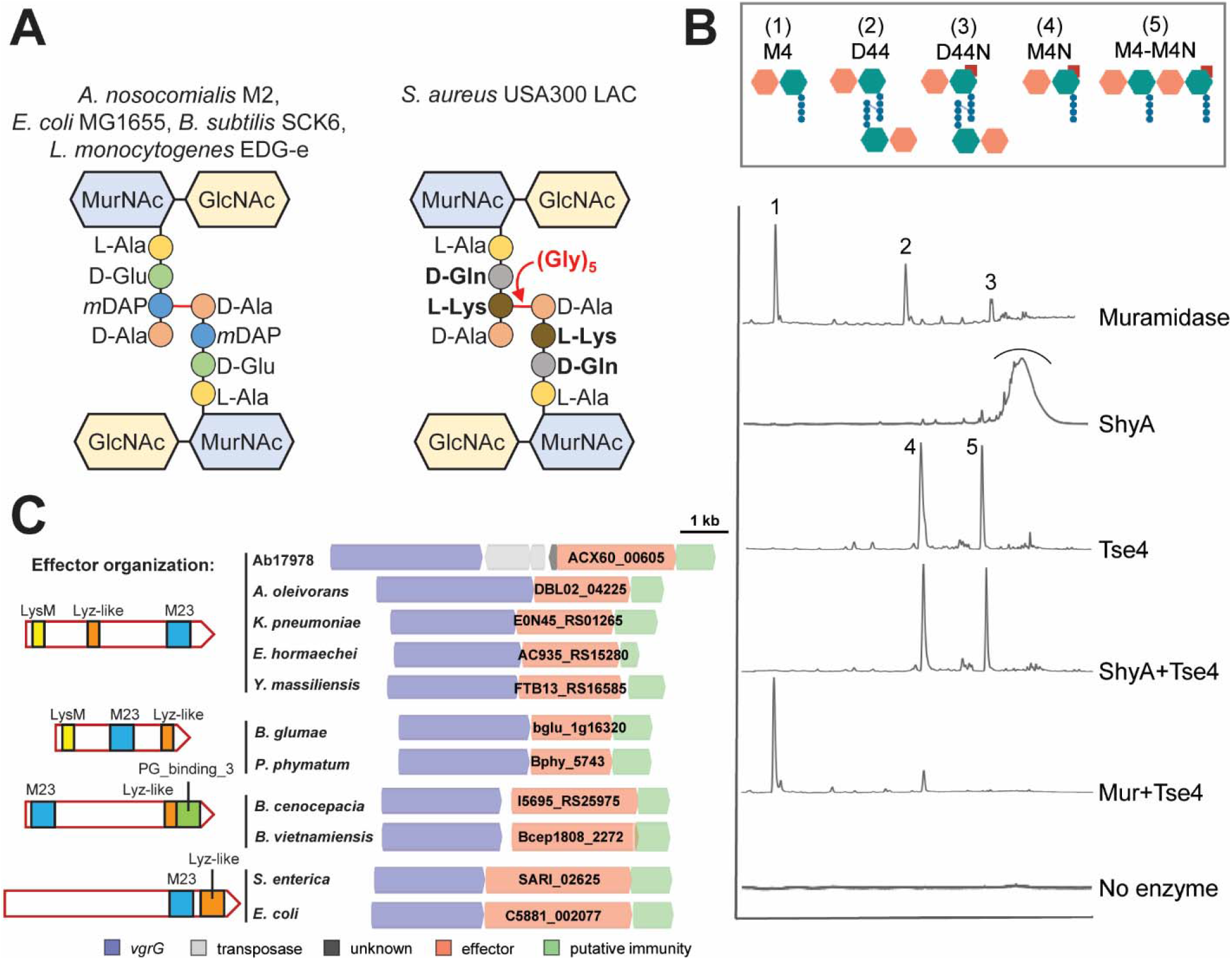
Tse4 is a bifunctional T6SS effector with modular architecture. **A.** PG composition of bacterial species susceptible to Tse4. **B.** Chromatograms of muropeptides released following PG treatment with the indicated enzymes. **C.** Domain architecture of various Tse4-like T6SS effectors. Effector locus tags are indicated.

To determine whether Tse4 possesses lytic transglycosylase and endopeptidase activities, we performed *in vitro* enzymatic assays followed by analysis of the solubilized muropeptides (PG fragments) by Ultra Performance Liquid Chromatography and Mass Spectrometry (UPLC-MS). Muramidase digestion of the PG produces a profile where the main peaks (muropeptides) are the monomeric disaccharide tetrapeptide (M4) and its dimeric (crosslinked) form (D44 and D44N) (Fig. 5B, S9). Subsequent addition of Tse4 converts the dimers to monomers (M4 and M4N), thereby confirming its endopeptidase activity. To test the putative lytic transglycosylase activity of Tse4, we incubated the PG first with the endopeptidase ShyA, which breaks the crosslinks to produce PG chains of diverse length and always ending with anhydro-M4 (M4N). Subsequent addition of Tse4 further processes the chains to shorter ones of 1 and 2 disaccharides long (M4N and M4-M4N) (Fig. 5B, S9), thereby confirming the lytic transglycosylase activity. As Tse4 cannot further break M4-M4N, we conclude that the predicted Lyz-like domain has endolytic transglycosylase activity. These results were recapitulated by digesting PG with Tse4 alone (Fig. 5B, S9). The identity of M4-M4N was confirmed both by MS and by the observation that this chain can be cleaved by muramidase into M4 and M4N (Fig. 5B, Table S1). Together, these data indicate that Tse4 is, to our knowledge, the first identified bifunctional T6SS effector.

A bioinformatics analysis identified orthologues of Tse4 in various *Acinetobacter*, *Klebsiella*, *Yersinia* and *Enterobacter* strains (Fig. 5C). Furthermore, we identified a series of T6SS effectors displaying a modular organization, in which the endopeptidase, lytic transglycosylase and PG-binding domain are linked in different arrangements (Fig. 5C). For example, a putative T6SS effector of *Burkholderia* spp. contains the same three domains but in the reverse order. Additional related effectors were identified in *E. coli* and *Salmonella enterica*. Thus, Tse4 belongs to a novel broadly distributed class of T6SS effectors that share a similar modular architecture. These findings suggest that Tse4-like effectors are important mediators of antagonistic bacterial interactions. In general, the modular arrangement of these newly discovered T6SS effectors resemble the organization of phage endolysins. Endolysins appear to have acquired multiple activities to enable host diversification and facilitate adaptation to specific growth phase- or strain-specific cell wall modifications(*16*). It is tempting to speculate that multicatalytic T6SS effectors evolved to provide a fitness advantage against diverse bacterial competitors. We propose that the remarkably broad target range and high potency of Tse4 is due to its dual catalytic activity, which might act synergistically to cause localized, lethal damage in the prey cell wall and/or to adjust to the variable PG chemistries of diverse prey.

Early work in *V. cholerae* and *P. aeruginosa* suggested that the T6SS targets Gram-negative but not Gram-positive bacteria(*17*, *18*). It was proposed that the PG of Gram-positive bacteria is too thick to allow the T6SS machinery to deliver toxic effectors at effective concentrations. However, recent work demonstrated that the T6SSs of *Serratia marcescens* and *Klebsiella pneumoniae* mediate fungal cell death(*19*, *20*), indicating that T6SS machines are able to penetrate and deliver toxic effectors across the fungal cell wall (>110nm), which is thicker than the cell wall of Gram-positive bacteria (<80nm). In this work, we show that Ab17978 employs its bifunctional T6SS effector Tse4 to kill various Gram-positive bacteria, including *B. subtilis*, *L. monocytogenes*, and *S. aureus*.

The outcome of interbacterial interactions is impacted by the environmental conditions in which these interactions occur(*9*). Here, we show that by secreting D-Lys, Ab17978 modifies its microenvironment to potentiate Tse4 activity and increase T6SS-dependent killing of Gram-negative and -positive prey. Our data suggests that the synergistic effect of D-Lys on Tse4 lethality is likely due to alkalization of the predator-prey zone of contact. Consistently, we found that Tse4 has enhanced activity under alkaline conditions. Thus, our work establishes a novel class of bifunctional, modularly organized, T6SS effectors effective against Gram-positive and -negative bacteria, highlights the role of D-amino acids in modulating the microenvironment, and redefines the function of the T6SS to include the warfare between Gram-negative and Gram-positive bacteria. Future work will be necessary to better understand the full scope of T6SS-mediated killing of Gram-negative and -positive bacteria and elucidate the effect of these interactions in shaping the composition of bacterial communities in the context of the human microbiota and polymicrobial infections.

## Acknowledgements

This work was supported by the National Institutes of Health grant 1R01AI125363-01 to M.F.F. Work in the Cava lab was supported by the Swedish Research Council (VR), the Knut and Alice Wallenberg Foundation (KAW), the Laboratory of Molecular Infection Medicine Sweden (MIMS), and the Kempe Foundation. We thank Dr. Ichiro Matsumura (Emory University) for providing *B. subtilis* SCK6 harboring pBAV1k-T5-gfp plasmid. We thank Dr. Laura Alvarez for providing ShyA. We thank Dr. Clay Jackson-Litteken for critical reading of the manuscript.

## Author Contributions

Conceived and designed the experiments: N.H.L., V.P., F.C. and M.F.F. Performed the experiments: N.H.L., V.P. and J.L. Analyzed the data: N.H.L., V.P., F.C. and M.F.F. Wrote the paper: N.H.L., J.L., and M.F.F.

## Supplementary Materials

### Materials and Methods

#### Bacterial strains and growth conditions

Bacterial strains used in this study are listed in Supplementary Table S2. Unless otherwise noted, strains were grown in lysogeny broth (LB) liquid medium at 37◻°C with shaking (200◻rpm). The antibiotics kanamycin (25 or 50◻μg/ml), gentamicin (20 μg/ml), chloramphenicol (15◻μg/ml) and carbenicillin (100 μg/ml) were added when necessary.

#### Construction of *A. baumannii* mutant strains

The primers used in this study are listed in Supplementary Table S3. Ab17978*Δtse4*, *ΔracK*, UPAB1*ΔtssM* and UPAB1*racK*+*ΔtssM* deletion mutants were constructed as described previously(*1*). Briefly, an antibiotic resistance cassette was amplified with primers pair P1 & P2 (Integrated DNA Technologies) with homology to the flanking regions of the targeted gene with additional 3′ 20 nucleotides of homology to the FRT site-flanked kanamycin resistance cassette from plasmid pKD4. This PCR product was electroporated into competent Ab17978, Ab17978*Δ123* unmarked mutant, UPAB1 or UPAB1*racK+* carrying plasmid pAT04, which expresses the RecAB recombinase. Mutants were selected on 10◻μg/ml kanamycin, and integration of the resistance marker was confirmed by PCR. To remove the kanamycin resistance cassette, electrocompetent mutants were transformed with plasmid pAT03, which expresses the FLP recombinase.

The *tsi4* gene was PCR amplified from Ab17978 and was cloned into vector pWH1266-promlac by Gibson Assembly (Hi-Fi DNA Assembly mix, NEB). The pWH1266-promlac-Tsi4_Ab17978 vector was electroporated into M2*ΔtssB* and selected on 15 μg/ml tetracycline, creating the M2ΔtssB::*tsi4* strain.

#### DAO enzymatic assay

3 mL of bacterial cultures grown for 20 h were centrifuged at 10,000 × g and 0.5 mL of supernatants were collected. The supernatants were deproteinized using Amicon Ultra-0.5 mL Centrifugal Filters with 3,000 MWCO. The flow through samples were then incubated with 0.4 mg/ml of D-amino acid oxidase DAO (Sigma-Aldrich) for 1 h at 37°C. Hydrogen peroxide co-product was then measured using HyPerBlu chemiluminescent detection kit (Lumigen), according to the manufacturer’s instructions.

#### Hcp secretion and Western blotting

Ab17978 overnight cultures were back-diluted in fresh LB medium to an OD600 of 0.025 and grown at 37°C with shaking until they reached an OD600 of 0.4 to 0.7. The cells were then pelleted by centrifugation. The cells were resuspended in Laemmli buffer to a final OD600 of 10. Supernatant proteins were subsequently precipitated with trichloroacetic acid, as previously described(*2*). Optical density-normalized volumes of whole cells or supernatants were loaded onto 15% SDS-PAGE gels for separation, transferred to a nitrocellulose membrane, and probed with polyclonal rabbit anti-Hcp (1:1,000) (39), polyclonal rabbit anti-6×His (1:2,000; Invitrogen, Waltham, MA) or monoclonal mouse anti-RNA polymerase (1:2,600; BioLegend, San Diego, CA). Western blots were then probed with IRDye-conjugated anti-mouse and anti-rabbit secondary antibodies (both at 1:15,000; LI-COR Biosciences, Lincoln, NE) and visualized with an Odyssey CLx imaging system (LI-COR Biosciences).

Detection of Hcp in solid growth was done by colony blot(*3*). 10 μL of overnight cultures normalized to OD600 of 1.0 were spotted onto 0.4 μm nitrocellulose membrane on top of an LB-agar (3%) plate. After 4 h of incubation at 37◻C, the membrane was washed and blotted as described above. Western-blot signals were quantified using LI-COR Image Studio software.

#### Protein expression and purification

The *tse4* gene was PCR amplified from Ab17978 and cloned into vector pET22b+ void of the pelB sequence using Gibson Assembly (Hi-Fi DNA Assembly mix, NEB). The pET22b+_tse4 vector was electroporated into *E. coli* Rosetta II (Invitrogen) and selected on 100◻μg/ml carbenicillin. The bacteria were grown in auto-inducible media(*4*) for 48 h at 20°C. Cells were harvested by centrifugation, resuspended in resuspension buffer (20 mM Tris-HCl pH 7.5, 300 mM NaCl, 10% glycerol, 10 μM b-ME, 0.1% Triton X100, 30 mM imidazole) and lysed by two passages through a cell disruptor at 35,000 psi. (Constant System ltd., Kennesaw, GA). Cell lysates were clarified by centrifugation at 10,000 rpm for 10 min, then passed over a nickel-nitrilotriacetic acid-agarose (Ni-NTA) column (Gold Bio, St. Louis, MO). The column was washed with resuspension buffer containing 60 mM imidazole, and Tse4 was eluted in resuspension buffer containing 150 mM imidazole. Purified Tse4 was buffer exchanged using Sephadex G25 PD10 column and stored in storing buffer (20 mM Tris-HCl pH 7.5, 150 mM NaCl, 10% glycerol) at −80°C.

#### Interbacterial competition assay

Interbacterial competition assays in solid media were performed as previously described(*5*). Briefly, predator and prey overnight cultures were pelleted, washed in fresh LB, and resuspended at an OD600 of 1.0. The cultures were mixed at a predator:prey ratio of 1:5 (Ab17978:M2*ΔtssB* or UPAB1:M2*ΔtssB*), 1:20 (Ab17978:MG1655), 1:10 (UPAB1:MG1655), 20:1 (Ab17978:JH642, Ab17978:SCK6, Ab17978:USA300 LAC or Ab17978:EDG-e), and 10 μl drops were spotted onto an LB-agar (3%) plate. To test the effect of different amino acids (D-Lys, D-Arg, D-Ala, D-Met, L-Lys or L-Arg (Sigma-Aldrich)) on Ab17978 T6SS-dependent killing capability, LB-agar plates were supplemented with each amino acid to a final concentration of 1, 5, 10 or 20 mM. In the pH-controlled killing assay, LB-agar plates were supplemented with either 100 mM MOPS pH 6.8 or 20 mM HEPES pH 8.0. After 4 h at 37°C (or 5 h for *L. monocytogenes* EDG-e prey), the spots were harvested, resuspended in 0.7 mL of LB broth, serially diluted and plated on LB-agar plates supplemented with appropriate antibiotics to determine the CFU of surviving prey (Table S2). CFU of surviving Ab17978 predator was also enumerated on chloramphenicol LB-agar plates.

To assess non-contact killing, we performed an interbacterial competition assay in liquid media. Briefly, overnight cultures of predators and prey normalized to OD600 of 1.0 were added at the same ratios used in the competition assay on solid media. Then, the co-cultures were concentrated 10 times by centrifugation and incubated for 4 h at 37◻C. Finally, 0.7 ml of LB broth was added to 40 μL of co-culture, serially diluted and plated on LB-agar plates supplemented with appropriate antibiotics to determine the CFU of surviving prey.

#### Bacterial cell lysis assay

*B. subtilis* SCK6 was grown to an OD600 of ~1.0. Cells were harvested and washed in reaction buffer (20 mM TrisHCl pH 8.0, 30 mM NaCl). No enzyme (control), lysozyme or Tse4 were added to final concentration of 7.5 μM. *B. subtilis* was then serially diluted and surviving CFUs were enumerated at time = 0 (pre-addition of enzymes), 1 h and 2 h (post-addition).

#### PG isolation, Remazol Brilliant Blue (RBB)-labeling, Tse4 *in vitro* reactions and UPLC analysis

PG was isolated from a stationary phase Ab17978 culture grown in LB, as previously described(*5*). Cells were collected by centrifugation for 15 min at 4°C and 7000 × g and resuspended in 25 mM phosphate buffer pH 6 (PB) to a final concentration of 0.2 mg/ml. Cell lysis was achieved by adding the cell suspension dropwise to an equal volume of boiling 8% (w/v) SDS under vigorous stirring and boiling the sample for an additional 30 min. After cooling to room temperature, the crude PG was collected by ultracentrifugation for 30 min at 110,000 × g at 25°C. Pellets were then washed several times with PB to remove SDS, treated with α-amylase (1 mg/ml) for 1 h at 37°C, and treated with Pronase E (2 mg/ml) overnight at 60°C. The reaction was stopped by boiling the sample with SDS, as described before, and residual SDS was removed by washing several times with PB.

RBB labeling of isolated PG was carried out as described previously(*6*, *7*). First, PG was resuspended in 0.02 M RBB 0.25 M NaOH to 12.5 mg/ml and incubated overnight at 37°C with agitation. The solution was then neutralized with HCl and RBB-labeled PG was pelleted by ultracentrifugation. Excess RBB was removed by repeatedly washing with water until no RBB was detected in the soluble fraction. RBB-labeled PG was then resuspended in 10% glycerol to 400 mg of wet pellet per mL and stored at −20°C. 40 mg/ml of RBB-dyed sacculi from Ab17978 were subjected to overnight reaction at 37°C with 7.5 μM of Tse4, lysozyme or no added enzyme (control reaction) in the reaction buffer (20 mM Tris-HCl at different pH, ranging from 6.8 to 10.5, 30 mM NaCl, 0.1 mM DTT, 0.1% Triton X100, 5 mM ZnCl_2_ and 7.5 mM EDTA). The resulting reactions were centrifuged at 14,000 × g to remove insoluble undigested RBB-dyed PG. The RBB-dye released in the reaction supernatant was quantified by absorbance at 585 nm.

Sacculi from *Vibrio cholerae* stationary phase cells were isolated as previously described(*8*). In short, PG sacculi were prepared by boiling bacteria in SDS (5% v/v). SDS was removed by several washes using ultracentrifugation, and the insoluble material was resuspended in MilliQ water. These sacculi were then subjected to overnight reactions with Tse4, ShyA and/or muramidase at 37°C, using an established protocol(*9*). For ShyA and Tse4, the *in vitro* reactions were performed in 100 mM Tris-HCl pH 8 containing 7.5 μM final concentration of enzyme. Muramidase reactions were carried out in MilliQ water or, when combined with Tse4, in 100 mM Tris-HCl pH 8. All assays were stopped by incubating the reaction at 100°C for 5 minutes. Soluble muropeptides were separated by Ultra Performance Liquid Chromatography (UPLC, Waters) and identified using a MALDI-TOF MS system (Waters).

#### Statistical analysis

All statistical analyses were performed using GraphPad Prism 8.0 (GraphPad Software Inc., La Jolla, CA). For all datasets, Student’s unpaired *t* tests were used.

---

**Fig. S1:**
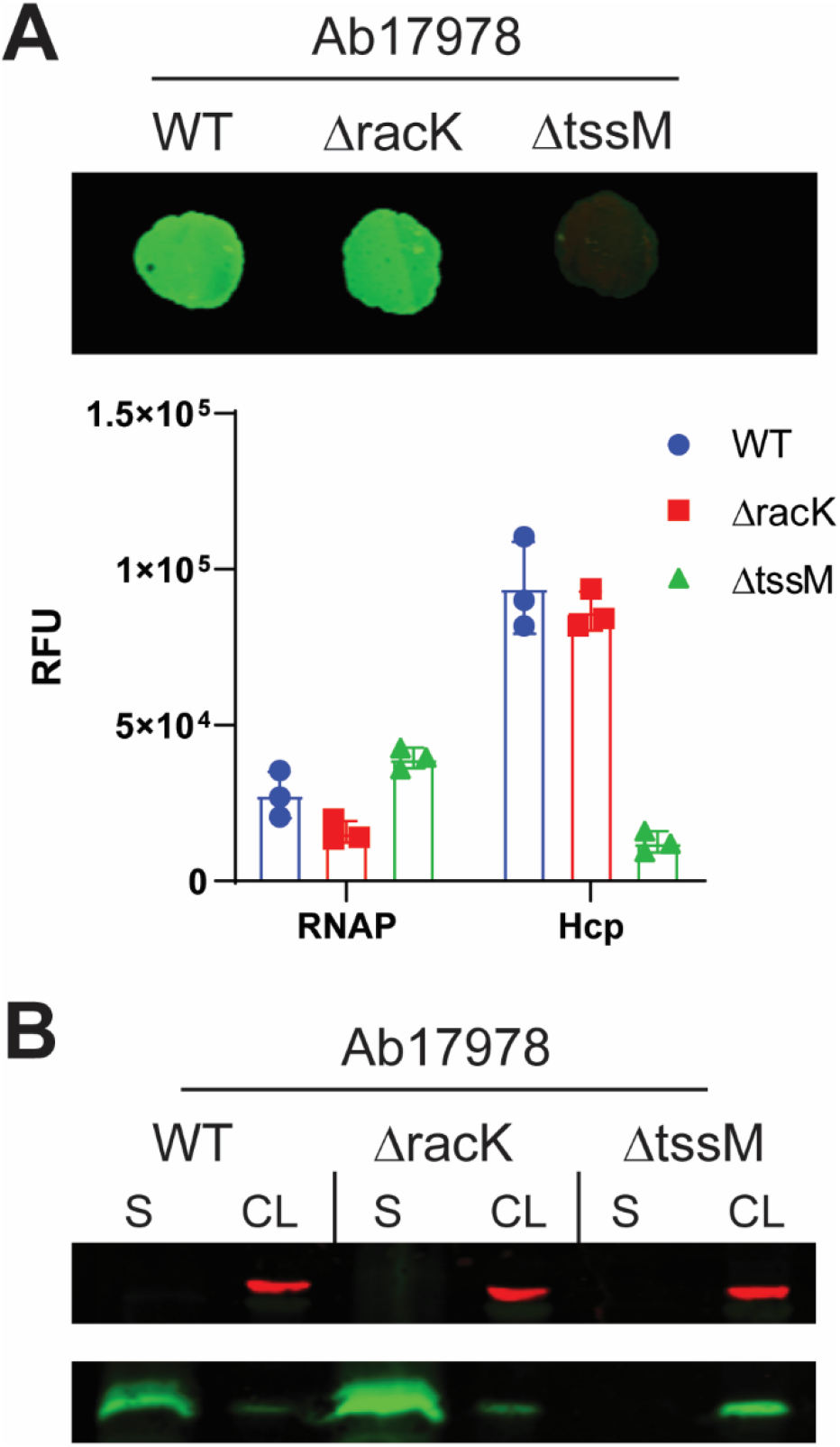
Deletion of *racK* does not impair Hcp secretion in Ab17978. **A.** (top) Representative colony blot of OD-normalized cell cultures of the indicated strains probing for Hcp and RNA polymerase (RNAP, lysis control). The cells were spotted on LB agar plates and allowed to grow for 4 h to replicate the killing assay conditions. A high signal of Hcp is indicative of T6SS activity. (bottom) Quantification of three independent colony blot experiments ± SD. **B.** Western blot of OD-normalized supernatant (S) and cell lysate (CL) fractions from the indicated Ab17978 strains probing for Hcp expression and secretion (green). RNAP is used as a lysis and loading control (red).

**Fig. S2:**
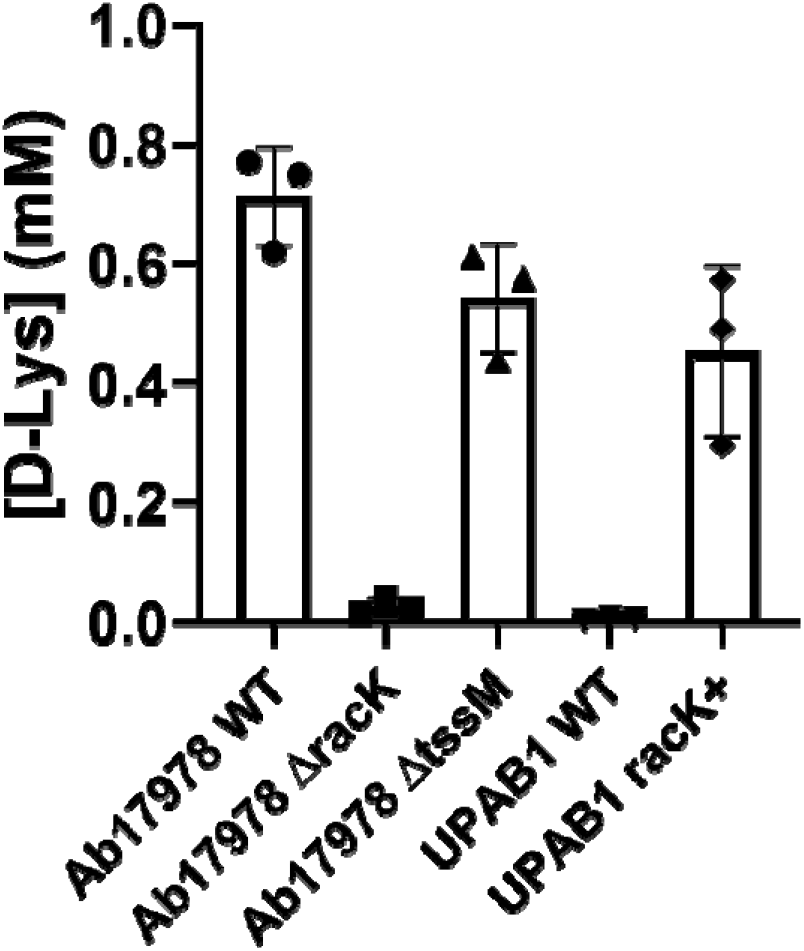
Secretion of D-Lys is independent of T6SS activity. Quantification of D-Lys present in supernatant fractions of the indicated strains using the DAO enzymatic assay. Data represent means ± SD of three biological replicates.

**Fig. S3:**
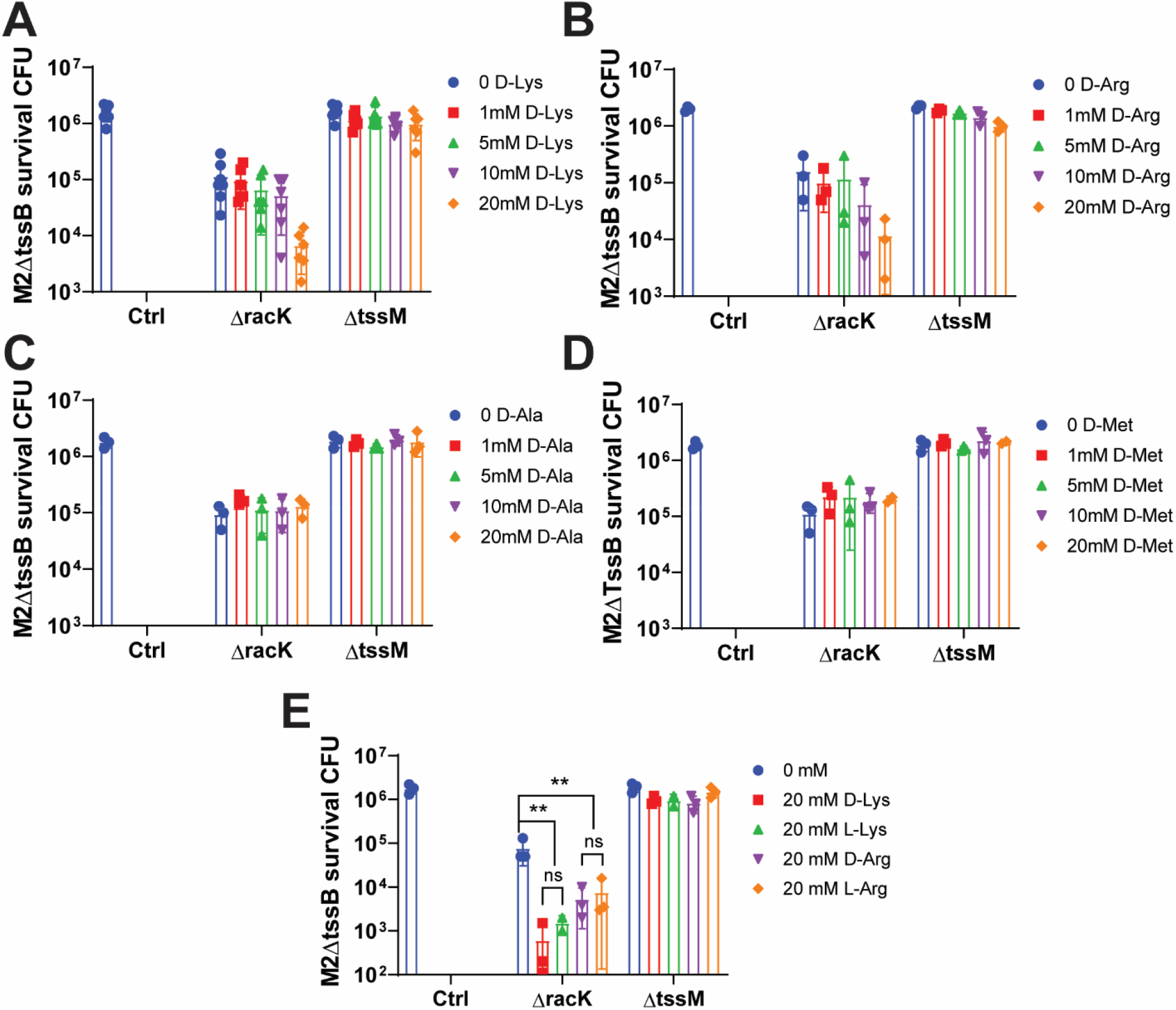
Extracellular L/D-Lys and −Arg enhance bacterial killing by Ab17978 *ΔracK*. Competition assay using Ab17978 *ΔracK* or *ΔtssM* as predators and M2*ΔtssB* as prey in LB-agar containing increasing amounts of (**A**) D-Lys, (**B**) D-Arg, (**C**) D-Ala, (**D**) D-Met, or (**E**) 20 mM of D-Lys, L-Lys, D-Arg or L-Arg. Control (Ctrl) indicates CFU of M2*ΔtssB* growing in LB-agar alone. Bar graphs represent means of prey survival CFU after 4 h of co-incubation of at least three biological replicates.

**Fig. S4:**
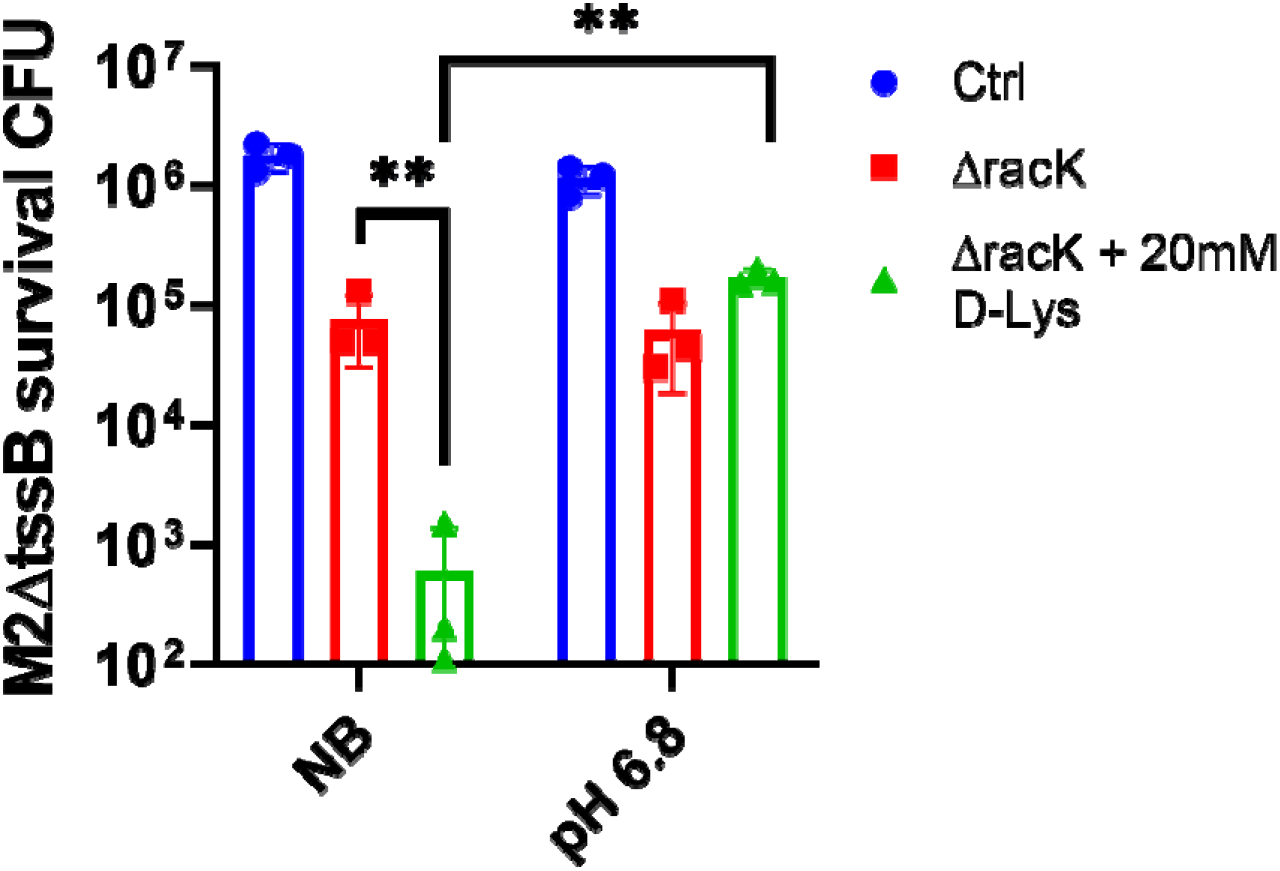
Buffering at neutral pH prevents the synergistic effect of D-Lys on bacterial killing. Competition assay using Ab17978*ΔracK* as predator and M2*ΔtssB* as prey in LB-agar with or without 20 mM D-Lys in either non-buffered (NB) media or media buffered at pH 6.8. Bar graphs represent means of prey survival CFU after 4 h of co-incubation of three biological replicates.

**Fig. S5:**
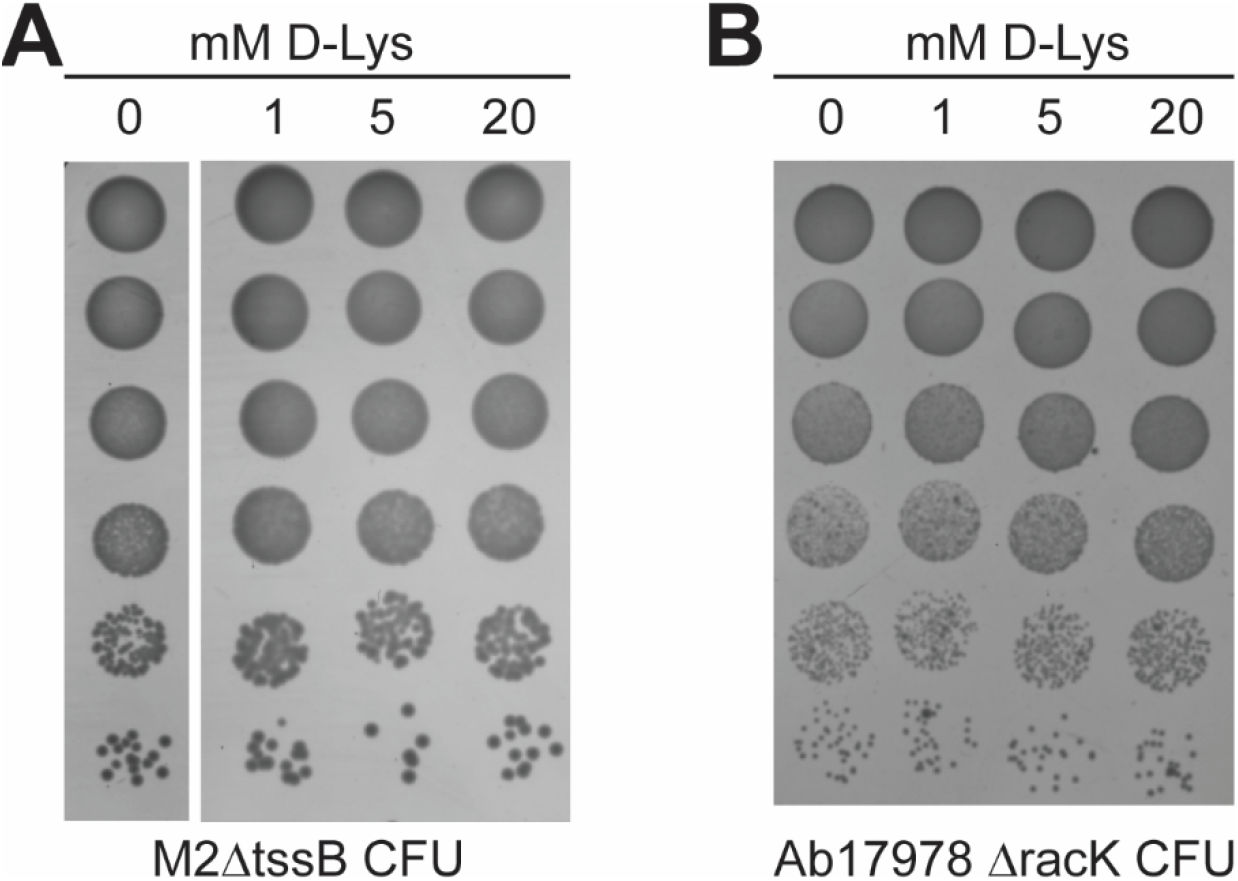
D-Lys does not affect the growth of M2 *ΔtssB* or Ab17978 *ΔracK*. Images showing survival CFU of (**A**) M2*ΔtssB* or **(B**) Ab17978*ΔracK* following a 4-h incubation in LB-agar supplemented with 0, 1, 5 or 20 mM of D-Lys on solid LB media. These conditions are identical to those used for competition assays.

**Fig. S6:**
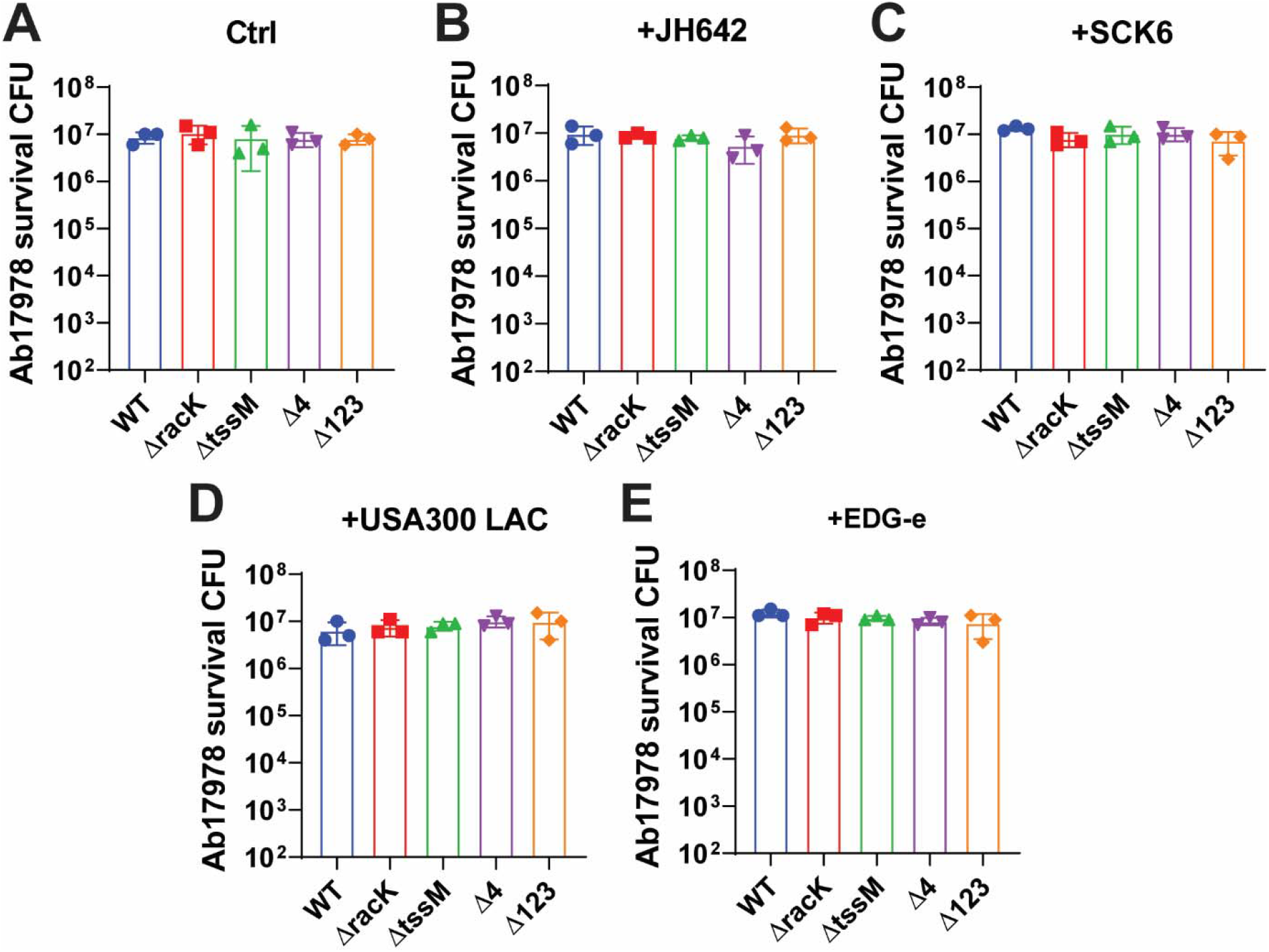
Ab17978 growth is unaffected by co-incubation with various Gram-positive bacteria in solid media. Bar graphs represent means of predator survival CFU after 4 h of co-incubation with the indicated prey strains (three biological replicates). The samples were obtained during the competition assays shown in Fig. 4.

**Fig. S7:**
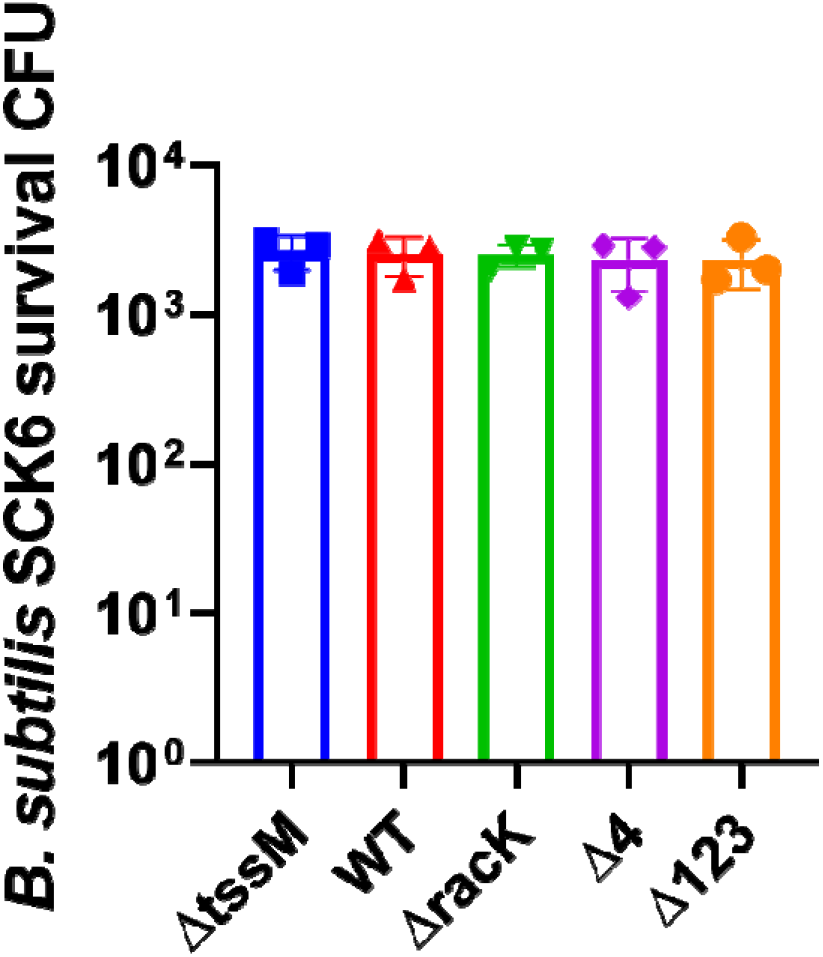
Ab17978 lacks bactericidal activity against *B. subtilis* SCK6 when co-incubated in liquid media. Interbacterial competition assay in liquid media using Ab17978 WT, *ΔtssM*, *ΔracK*, *Δ4* or *Δ123* as predators and *B. subtilis* SCK6 as prey. Bar graphs represent means of prey survival CFU after 4 h of co-incubation (three biological replicates).

**Fig. S8:**
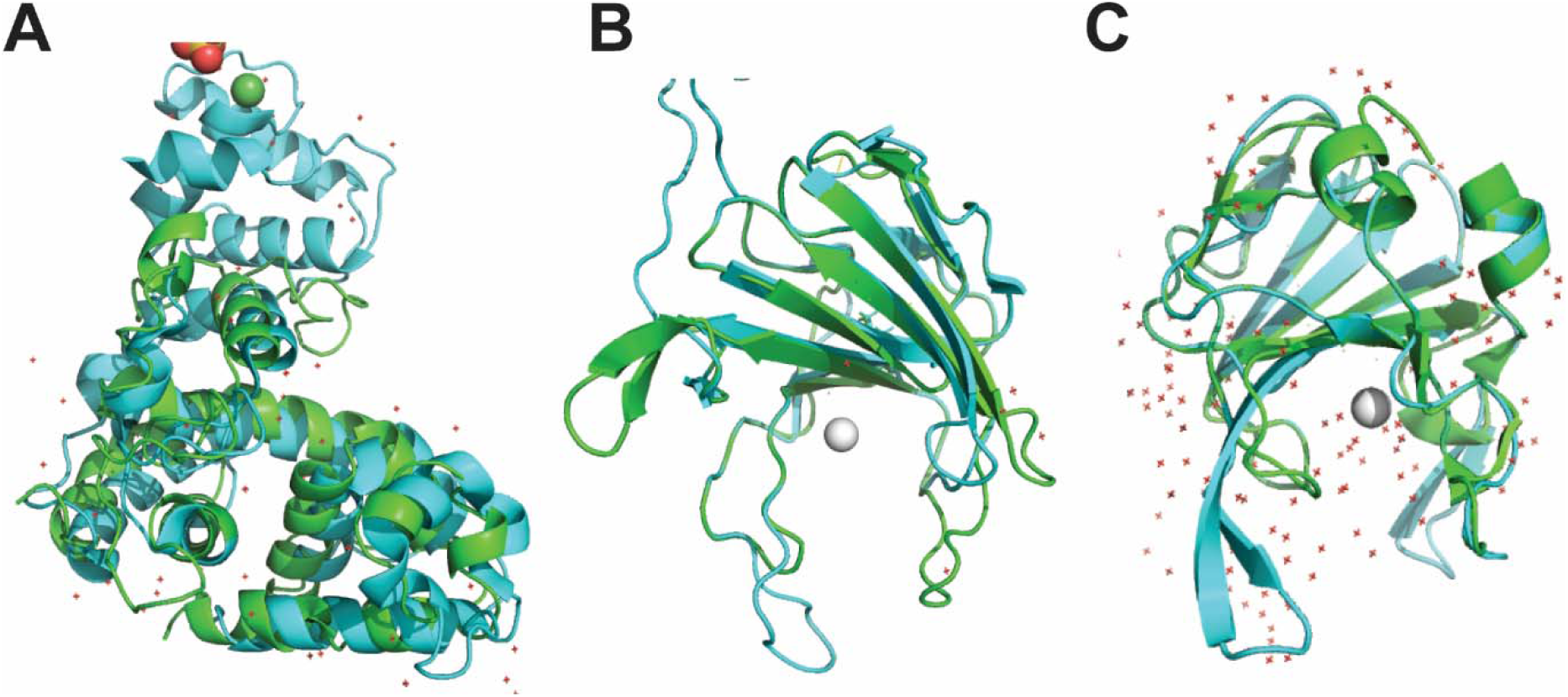
Tse4 is predicted to be a bifunctional effector with lysozyme and peptidase domains. (**A**) Tse4 (res. 242-447) modeled to lytic transglycosylase gp144 of bacteriophage phiKZ (PDB: 3BKH). Tse4 (res. 634-798) modeled to (**B**) the DD-endopeptidase ShyA of *V. cholerae* (PDB:6U2A) and (**C**) to lysostaphin LytM of *S. aureus* (PDB: 2B44). Structural models of Tse4 are shown in green; templates are shown in cyan.

**Fig. S9:**
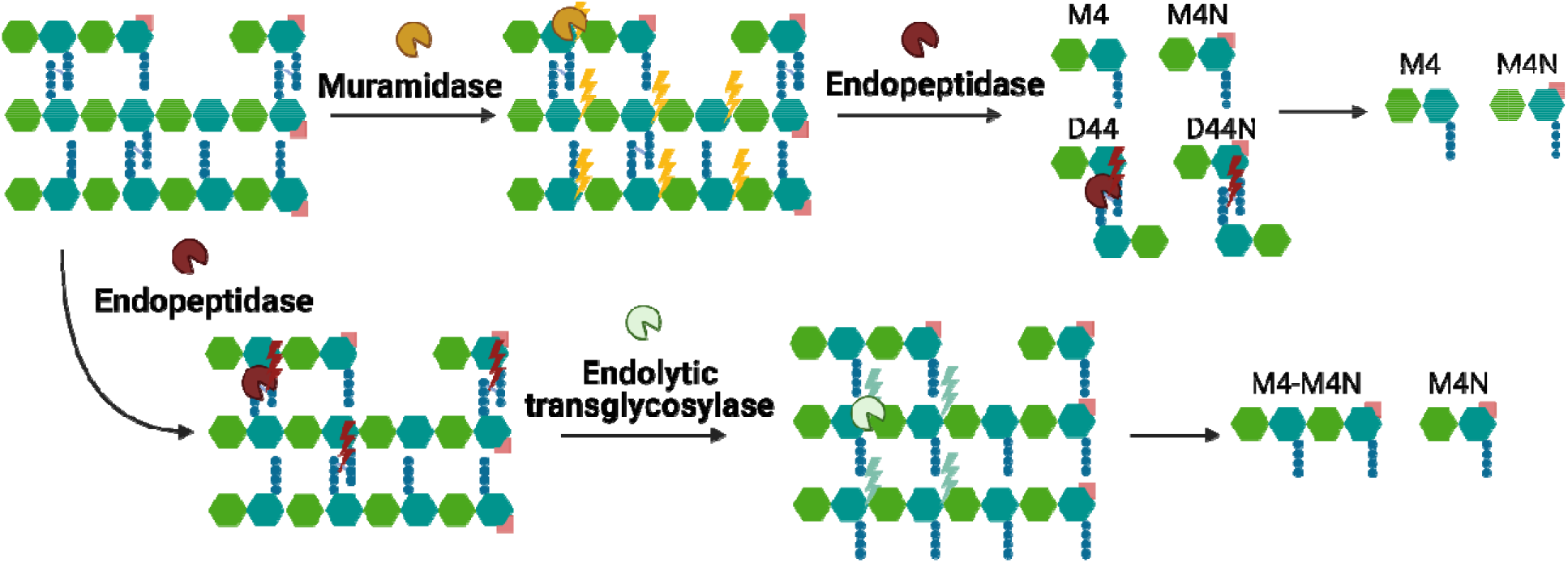
Schematic of the experimental design used to determine the bifunctionality of Tse4. PG is a heteromeric structure consisting of crosslinked and uncrosslinked muropeptide chains with an anhydromuropeptide marking the end of the glycosidic chain. (top) Muramidase cleaves the glycosidic link between MurNAc (blue) and GlcNAc (green), leading to the release of a mixture of crosslinked (D44) and uncrosslinked (M4) muropeptides, or their anhydro forms (D44N and M4N, respectively). Endopeptidases cleave the peptide stem. Thus, co-treatment of PG with muramidase+endopeptidase is expected to release exclusively uncrosslinked products (M4 and M4N). Like muramidases, endolytic transglycosylases cleave glycosidic bonds. However, unlike muramidases, endolytic transglycosylases form unique anhydromuropeptides. (bottom) Because endopeptidases do not cleave the glycan backbone of PG, endopeptidase digestion of the sacculi generates polymeric MurNAc-GlcNAc chains bearing non-crosslinked stems. In contrast, co-treatment of PG with endopeptidase+endolytic transglycosylase is expected to release large amounts of dimeric and monomeric anhydromuropeptides (M4-M4N and M4N, respectively).

**Table S1:** Muropeptide nomenclature and mass for chromatograms shown in Fig. 5.

| **(Peak)**<br>**Muropeptide** | **Predicted**<br>**mass** | **Observed**<br>**mass** |
| --- | --- | --- |
| (1) M4 | 942.4155 | 942.4232 |
| (2) D44 | 1865.8126 | 1865.8698 |
| (3) D44N | 1845.7864 | 1845.8116 |
| (4) M4N | 922.3893 | 922.39284 |
| (5) M4-M4N | 1843.7708 | 1843.7554 |

**Table S2:**
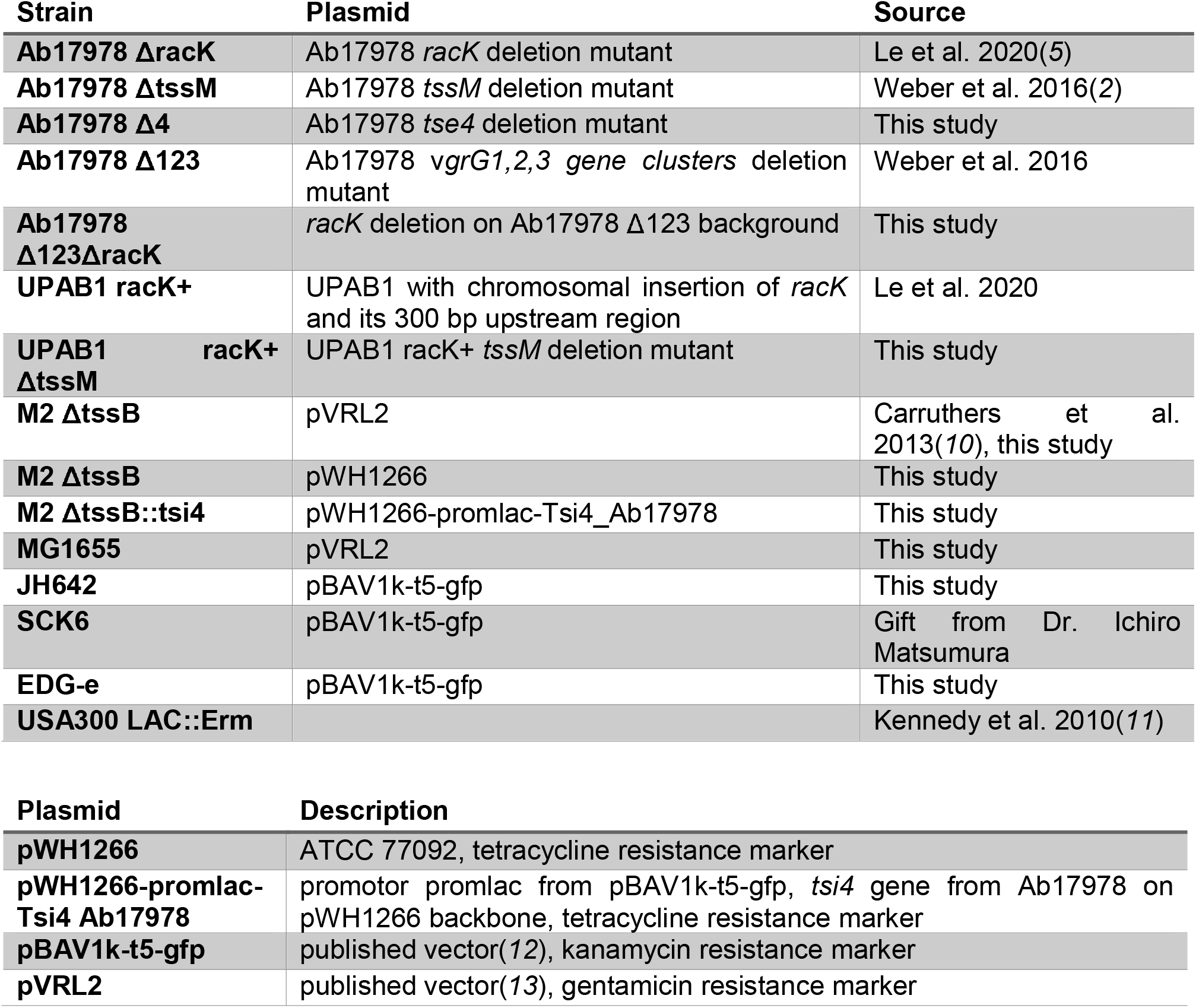
Bacterial strains and plasmids used in this study.

**Table S3:**
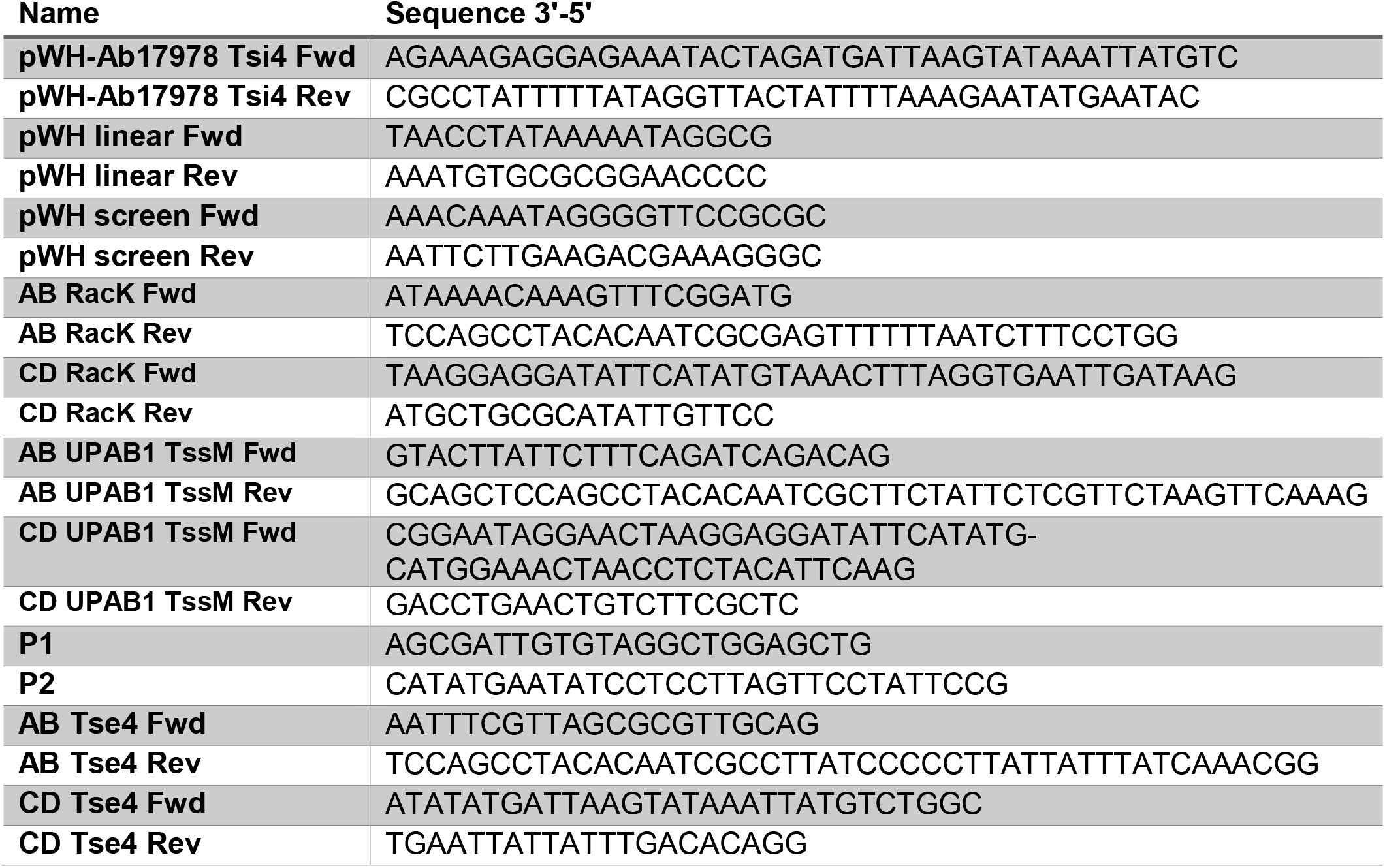
Primers used in this study.

## Bibliography

1. S. J. Hersch, K. Manera, T. G. Dong, Defending against the Type Six Secretion System: beyond Immunity Genes The bacterial type six secretion system (T6SS) delivers toxic effector proteins into neighboring cells (2020), doi:10.1016/j.celrep.2020.108259.

2. N. H. Le, K. Peters, A. Espaillat, J. R. Sheldon, J. Gray, G. Di Venanzio, J. Lopez, B. Djahanschiri, E. A. Mueller, S. W. Hennon, P. A. Levin, I. Ebersberger, E. P. Skaar, F. Cava, W. Vollmer, M. F. Feldman, Peptidoglycan editing provides immunity to Acinetobacter baumannii during bacterial warfare. Sci. Adv. 6, 5614–5636 (2020).

3. W. P. J. Smith, A. Vettiger, J. Winter, T. Ryser, L. E. Comstock, M. Basler, K. R. Foster, The evolution of the type VI secretion system as a disintegration weapon. PLoS Biol. 18, e3000720 (2020).

4. A. B. Russell, R. D. Hood, N. K. Bui, M. LeRoux, W. Vollmer, J. D. Mougous, Type VI secretion delivers bacteriolytic effectors to target cells. Nature. 475, 343–347 (2011).

5. J. C. Whitney, S. Chou, A. B. Russell, J. Biboy, T. E. Gardiner, M. A. Ferrin, M. Brittnacher, W. Vollmer, J. D. Mougous, Identification, structure, and function of a novel type VI secretion peptidoglycan glycoside hydrolase effector-immunity pair. J. Biol. Chem. 288, 26616–26624 (2013).

6. J. Ma, M. Sun, Z. Pan, C. Lu, H. Yao, Diverse toxic effectors are harbored by vgrG islands for interbacterial antagonism in type VI secretion system. Biochim. Biophys. Acta - Gen. Subj. 1862, 1635–1643 (2018).

7. A. B. Russell, P. Singh, M. Brittnacher, N. K. Bui, R. D. Hood, M. A. Carl, D. M. Agnello, S. Schwarz, D. R. Goodlett, W. Vollmer, J. D. Mougous, A widespread bacterial type VI secretion effector superfamily identified using a heuristic approach. Cell Host Microbe. 11, 538–549 (2012).

8. A. Aliashkevich, L. Alvarez, F. Cava, New insights into the mechanisms and biological roles of D-amino acids in complex eco-systems. Front. Microbiol. 9 (2018), p. 683.

9. K. D. LaCourse, S. B. Peterson, H. D. Kulasekara, M. C. Radey, J. Kim, J. D. Mougous, Conditional toxicity and synergy drive diversity among antibacterial effectors. Nat. Microbiol. 3, 440–446 (2018).

10. B. S. Weber, S. W. Hennon, M. S. Wright, N. E. Scott, V. de Berardinis, L. J. Foster, J. A. Ayala, M. D. Adams, M. F. Feldman, Genetic dissection of the type VI secretion system in Acinetobacter and identification of a novel peptidoglycan hydrolase, TagX, required for its biogenesis. MBio. 7(2016), doi:10.1128/mBio.01253-16.

11. F. Cava, M. A. de Pedro, H. Lam, B. M. Davis, M. K. Waldor, Distinct pathways for modification of the bacterial cell wall by non-canonical D-amino acids. EMBO J. 30, 3442–3453 (2011).

12. J. C. Whitney, C. M. Beck, Y. A. Goo, A. B. Russell, B. N. Harding, J. A. De Leon, D. A. Cunningham, B. Q. Tran, D. A. Low, D. R. Goodlett, C. S. Hayes, J. D. Mougous, Genetically distinct pathways guide effector export through the type VI secretion system. Mol. Microbiol. 92, 529–542 (2014).

13. W. Vollmer, D. Blanot, M. A. de Pedro, Peptidoglycan structure and architecture. FEMS Microbiol. Rev. 32, 149–167 (2008).

14. T. Dörr, F. Cava, H. Lam, B. M. Davis, M. K. Waldor, Substrate specificity of an elongation-specific peptidoglycan endopeptidase and its implications for cell wall architecture and growth of Vibrio cholerae. Mol. Microbiol. 89, 949–962 (2013).

15. M. Firczuk, A. Mucha, M. Bochtler, Crystal structures of active LytM. J. Mol. Biol. 354, 578–590 (2005).

16. H. Oliveira, L. D. R. Melo, S. B. Santos, F. L. Nobrega, E. C. Ferreira, N. Cerca, J. Azeredo, L. D. Kluskens, Molecular Aspects and Comparative Genomics of Bacteriophage Endolysins. J. Virol. 87, 4558–4570 (2013).

17. D. L. MacIntyre, S. T. Miyata, M. Kitaoka, S. Pukatzki, The Vibrio cholerae type VI secretion system displays antimicrobial properties. Proc. Natl. Acad. Sci. U. S. A. 107, 19520–19524 (2010).

18. S. Chou, N. K. Bui, A. B. Russell, K. W. Lexa, T. E. Gardiner, M. LeRoux, W. Vollmer, J. D. Mougous, Structure of a Peptidoglycan Amidase Effector Targeted to Gram-Negative Bacteria by the Type VI Secretion System. Cell Rep. 1, 656–664 (2012).

19. K. Trunk, J. Peltier, Y. C. Liu, B. D. Dill, L. Walker, N. A. R. Gow, M. J. R. Stark, J. Quinn, H. Strahl, M. Trost, S. J. Coulthurst, The type VI secretion system deploys antifungal effectors against microbial competitors. Nat. Microbiol. 3, 920–931 (2018).

20. D. Storey, A. McNally, M. Åstrand, J. P. G. Santos, I. Rodriguez-Escudero, B. Elmore, L. Palacios, H. Marshall, L. Hobley, M. Molina, V. J. Cid, T. A. Salminen, J. A. Bengoechea, Klebsiella pneumoniae type VI secretion system-mediated microbial competition is PhoPQ controlled and reactive oxygen species dependent. PLoS Pathog. 16(2020), doi:10.1371/journal.ppat.1007969.

## Additional references

1. A. T. Tucker, E. M. Nowicki, J. M. Boll, G. A. Knauf, N. C. Burdis, M. S. Trent, B. W. Davies, Defining Gene-Phenotype Relationships in Acinetobacter baumannii through One-Step Chromosomal Gene Inactivation. MBio. 5, e01313–14 (2014).

2. B. S. Weber, S. W. Hennon, M. S. Wright, N. E. Scott, V. de Berardinis, L. J. Foster, J. A. Ayala, M. D. Adams, M. F. Feldman, Genetic dissection of the type VI secretion system in Acinetobacter and identification of a novel peptidoglycan hydrolase, TagX, required for its biogenesis. MBio. 7(2016), doi:10.1128/mBio.01253-16.

3. B. S. Weber, P. M. Ly, M. F. Feldman, in Methods in Molecular Biology (Humana Press Inc., 2017; https://pubmed.ncbi.nlm.nih.gov/28667630/), vol. 1615, pp. 465–472.

4. F. W. Studier, Protein production by auto-induction in high-density shaking cultures (2005), doi:10.1016/j.pep.2005.01.016.

5. N. H. Le, K. Peters, A. Espaillat, J. R. Sheldon, J. Gray, G. Di Venanzio, J. Lopez, B. Djahanschiri, E. A. Mueller, S. W. Hennon, P. A. Levin, I. Ebersberger, E. P. Skaar, F. Cava, W. Vollmer, M. F. Feldman, Peptidoglycan editing provides immunity to Acinetobacter baumannii during bacterial warfare. Sci. Adv. 6, 5614–5636 (2020).

6. R. Zhou, S. Chen, P. Recsei, A dye release assay for determination of lysostaphin activity. Anal. Biochem. 171, 141–144 (1988).

7. T. Dörr, F. Cava, H. Lam, B. M. Davis, M. K. Waldor, Substrate specificity of an elongation-specific peptidoglycan endopeptidase and its implications for cell wall architecture and growth of Vibrio cholerae. Mol. Microbiol. 89, 949–962 (2013).

8. L. Alvarez, S. B. Hernandez, M. A. De Pedro, F. Cava, in Methods in Molecular Biology (Humana Press Inc., 2016; https://pubmed.ncbi.nlm.nih.gov/27311661/), vol. 1440, pp. 11–27.

9. L. Alvarez, S. B. Hernandez, M. A. De Pedro, F. Cava, in Methods in Molecular Biology (Humana Press Inc., 2016), vol. 1440, pp. 11–27.

10. M. D. Carruthers, P. A. Nicholson, E. N. Tracy, R. S. Munson, Acinetobacter baumannii Utilizes a Type VI Secretion System for Bacterial Competition. PLoS One. 8, e59388 (2013).

11. A. D. Kennedy, J. B. Wardenburg, D. J. Gardner, D. Long, A. R. Whitney, K. R. Braughton, O. Schneewind, F. R. DeLeo, Targeting of alpha-hemolysin by active or passive immunization decreases severity of USA300 skin infection in a mouse model. J. Infect. Dis. 202, 1050–1058 (2010).

12. A. V. Bryksin, I. Matsumura, Rational Design of a Plasmid Origin That Replicates Efficiently in Both Gram-Positive and Gram-Negative Bacteria. PLoS One. 5, e13244 (2010).

13. M. Lucidi, F. Runci, G. Rampioni, E. Frangipani, L. Leoni, P. Visca, New shuttle vectors for gene cloning and expression in multidrug-resistant Acinetobacter species. Antimicrob. Agents Chemother. 62(2018), doi:10.1128/AAC.02480-17.

